# Sniff invariant odor coding

**DOI:** 10.1101/174417

**Authors:** Roman Shusterman, Yevgeniy B. Sirotin, Matthew C. Smear, Yashar Ahmadian, Dmitry Rinberg

## Abstract

Sampling regulates stimulus intensity and temporal dynamics at the sense organ. Despite variations in sampling behavior, animals must make veridical perceptual judgments about external stimuli. In olfaction, odor sampling varies with respiration, which influences neural responses at the olfactory periphery. Nevertheless, rats were able to perform fine odor intensity judgments despite variations in sniff kinetics. To identify the features of neural activity supporting stable intensity perception, in awake mice we measured responses of Mitral/Tufted (MT) cells to different odors and concentrations across a range of sniff frequencies. Amplitude and latency of the MT cells’ responses vary with sniff duration. A fluid dynamics (FD) model based on odor concentration kinetics in the intranasal cavity can account for this variability. Eliminating sniff waveform dependence of MT cell responses using the FD model significantly improves concentration decoding. This suggests potential schemes for sniff waveform invariant odor concentration coding.

**Highlights:**

- Odor concentration discrimination does not depend on sniff frequency
- Amplitude and latency of MT cell responses vary with sniff frequency
- A fluid dynamic based model accounts for sniff dependent variability in the responses
- Transforming MT cell responses with this model achieves sniff invariant coding

## Introduction

Sensory systems acquire and process external stimuli. Active sensing allows an animal to control the acquisition of this information. The eyes target and scan objects of interest. The hands grasp and palpate objects. However, variation in sampling behavior introduces variability in the acquired signal. Sensory systems must incorporate or correct for this variability to achieve perceptual constancy.

In olfaction, odor sampling is controlled by sniffing, which determines the time course of odor stimulation (Wachowiak, 2011). In awake animals, sniff waveforms are variable. Thus, odor stimuli at a fixed concentration will result in varying stimuli at the olfactory epithelium. Nevertheless, observations from human psychophysics support the idea of independence of odor intensity perception on sniff airflow (Mainland et al., 2014; R. Teghtsoonian and M. Teghtsoonian, 1984; R. Teghtsoonian et al., 1978). Additional evidence comes from rodent behavioral experiments, which show that animals can discriminate odors equally well in a freely moving assay (fast sniffing) as in a head restrained one (slow breathing) (Wesson et al., 2009). However, it may be problematic to compare results from different behavioral assays.

Variation in sniffing has been shown to influence neural responses of early olfactory neurons. OSN activation is coupled to inhalation and significantly altered by changes in sampling frequency (Carey et al., 2008; Ghatpande and Reisert, 2011; Spors et al., 2006; Verhagen et al., 2007). Previous studies using artificial sniffing found that responses of mitral/tufted (MT) cells in the olfactory bulb (OB) of anesthetized rats depend on sniff rate and magnitude (Courtiol et al., 2011; Esclassan et al., 2012). Most MT cells are modulated by respiration at low and moderate sniff frequencies. At higher frequencies, response magnitude is reduced (Bathellier et al., 2008; Carey and Wachowiak, 2011) while response latency remains unchanged (Carey and Wachowiak, 2011). Recordings from awake animals show controversial results: Cury, et al, showed that responses do not depend on sniff frequency (Cury and Uchida, 2010), while our previous work showed MT cell response latencies scale with parameters of the sniff cycle (Shusterman et al., 2011). Given that MT cells’ responses vary with sniff parameters, how is stable odor perception achieved?

Various candidate odor representations with sniff frequency invariance have been proposed. Some studies have argued that spike trains of MT cells are locked to the phase of the sniff cycle (Chaput, 1986; Roux et al., 2006). In this scheme, which we call “phase” transformation, all spike trains were stretched or compressed to match the duration of an average sniff cycle. In our previous work (Shusterman et al., 2011), we proposed to do a more elaborate transformation by independently stretching and compressing the inhalation interval and the rest of the sniff interval – “two-interval phase” transformation. Another proposed representation scales the whole sniff interval based on the duration of the inhalation interval – “inhalation proportional” transformation (Arneodo et al., 2017). The purpose of all these transformations was to eliminate the sniff waveform odor response dependences, and find a sniff frequency invariant representation. However, the thorough comparison of these models, and quantification of how well they perform is still lacking.

In this study, we first tested whether sampling affects odor perception. In rats performing a two-alternative choice task, odor concentration discrimination does not vary with sampling frequency. To investigate which features of the neural code underlie this stable performance, we exploited the natural variability in the sniffing pattern of awake mice and recorded MT cells’ responses to different odorants and different concentrations of the same odor. We found that for different sniff patterns, MT cell responses during the early part of the sniff cycle are less variable than the responses during the later part. However, even early responses vary systematically with sniff duration.

Based on physical principles of airflow propagation in the nose, we proposed a simple fluid dynamics (FD) model. Transformation of MT cells responses according to the FD model explains the systematic variability in neural responses. We compared odor responses after transformation based on this model against 4 earlier proposed transformations and showed that FD transformation provided the best description of the data. Further, discriminant analysis showed that FD transformation improves concentration discrimination. Based on these observations, we propose a transformation of MT cell responses which is invariant to sniff waveform parameters, and discuss how such a representation can be read by higher brain areas.

## Results

### Odor concentration perception does not depend on sniff duration

To test whether odor intensity perception depends on sniff duration, we trained rats to classify odor concentrations in a two-alternative choice task (Fig. 1A, B). On each trial, we presented randomly one of eight odor concentrations. Rats were trained to go to the left water port for four higher concentrations and to go to the right water port for the four lower concentrations (Fig. 1A). Rats performed on average about 700-800 trials per session which results in ∼80-100 trials per each concentration. We estimated behavioral performance using psychometric functions (Fig. 1C) with three parameters: *μ* - the decision boundary between high and low concentrations; *σ* - the curve slope, which corresponds to odor concentration discriminability (sensitivity); and *λ* - lapse rate, the percentage of trials on which the animal guessed the answer (Wichmann and Hill, 2001a).

**Figure 1.**
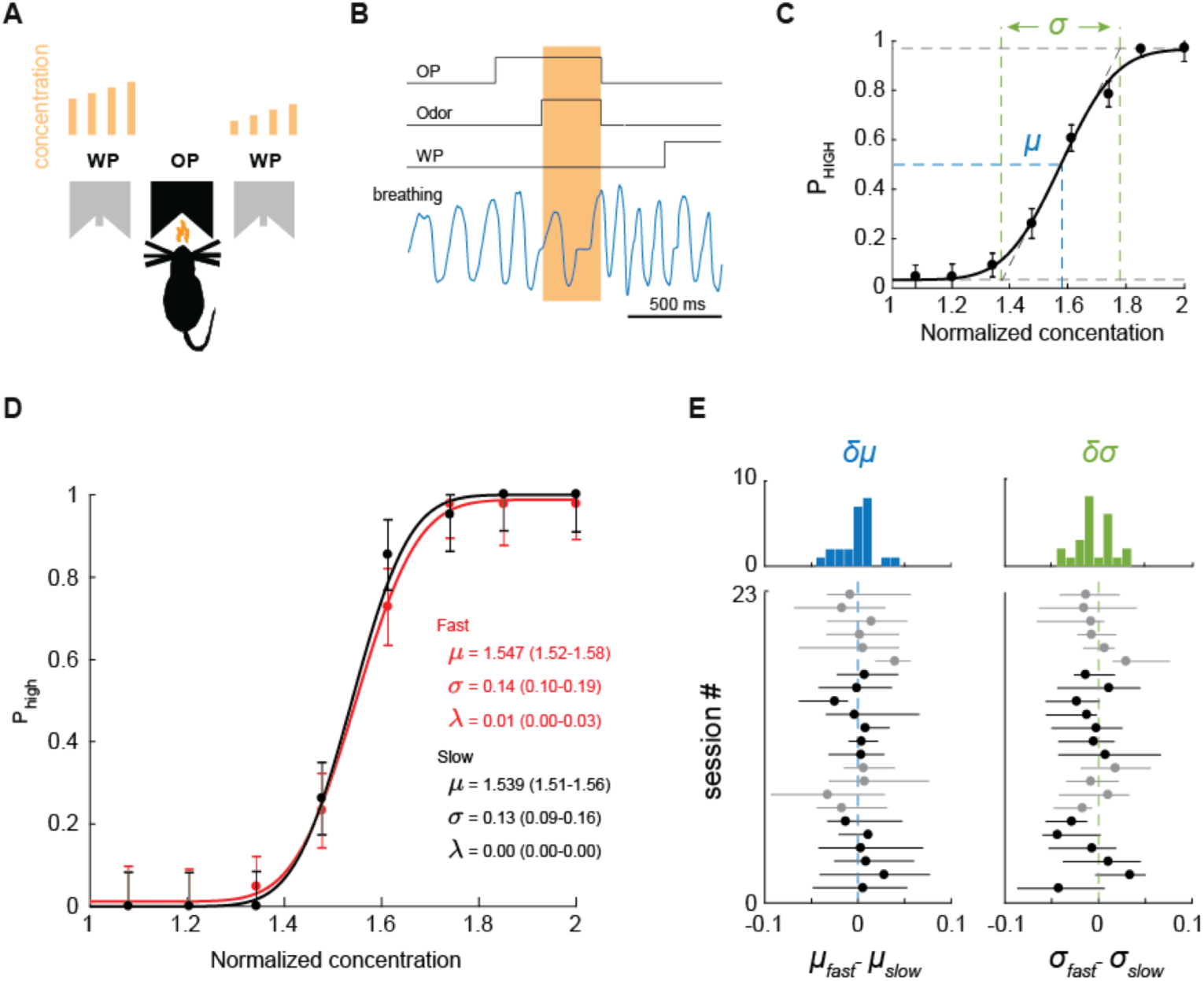
Behavioral discrimination of odor concentrations is sniff frequency invariant. **A.** Two-alternative choice odor discrimination task. Rats initiate trials by poking into an odor port (OP), which activates presentation of 1 out of 8 odor concentrations. After being exposed to one of 4 low/high concentrations, rats are rewarded for poking into right/left water port (WP). Light brown bars above the water ports represent the different concentrations the rat must discriminate. **B.** Task trial structure and pressure signal. Downward and upward deflections of the pressure signal reflect inhalation and exhalation, respectively. Light brown shading marks odor presentation time interval. **C.** Average concentration classification performance (n = 4 rats). Error bars denote standard deviations across all sessions for all rats. The continuous line is an error function fit to the data. **D.** Example performance during a single session for fast (black) and slow (red) sniff cycles. Each point denotes probability of going left (high concentration). Each inset is a sniff duration histogram for a given concentration. **E.** Differences in psychometric parameters between fast and slow sampling for decision boundary, *δμ = μ*_*fast*_ - μ_*slow*_, and sensitivity, *δσ = σ* _*fast*_ - σ_*slow*_: histograms for all sessions (top) and values for individual sessions (bottom). Lines are 95% confidence intervals assessed via bootstrap. Sessions from different animals are presented in alternating black and gray colors.

At the first sniff after odor exposure, rats exhibited a broad range of sniff durations (across all rats: median: 143 ms, range: 74-441 ms) (Fig. S1, inserts). To quantify the dependence of animal performance on sniff duration we split all trials into two groups: with fast/slow sniffs (sniff duration was shorter/longer than the median sniff duration for a given rat). We fit psychometric functions independently for each group of trials (Fig. 1D). The differences between groups for the decision boundaries *δ*_*μ*_=*μ*_*fast*_ - *μ*_*slow*_, and sensitivities *δ*_*σ*_=*σ*_*fast*_ - *σ*_*slow*_, did not statistically differ from zero (*δ*_*μ*_: p=0.77, *δ*_*σ*_: p=0.18, Wilcoxon signed rank test). These results suggest that neither concentration sensitivity, *σ,* nor biases in perceived concentration, *μ*, depend on the sniff duration. Therefore, behavioral performance in this concentration discrimination task does not depend on sniff duration.

## Odor responses vary with sniff duration

To investigate which features of neural code are invariant to variations in sniff pattern, we exploited the natural variability in the sniffing pattern of awake animals. We recorded MT cell responses (13 mice, 200 cells) to different odorants and different concentrations of the same odor in head-restrained mice (Fig. 2A and Methods). Sniffing was monitored via an intranasal pressure cannula (Fig. 2A). Responses of each MT cell were recorded typically to 5-10 odorants, making in total 1409 cell-odor pairs. Out of 200 MT cells, 134 cells were recorded in sessions with a presentation of multiple odor concentrations (548 unit–odor– concentration triplets). Out of all responses, 28% demonstrated initially excitatory activity and 28% initially inhibitory (Fig. 2B) (see Methods).

**Figure 2.**
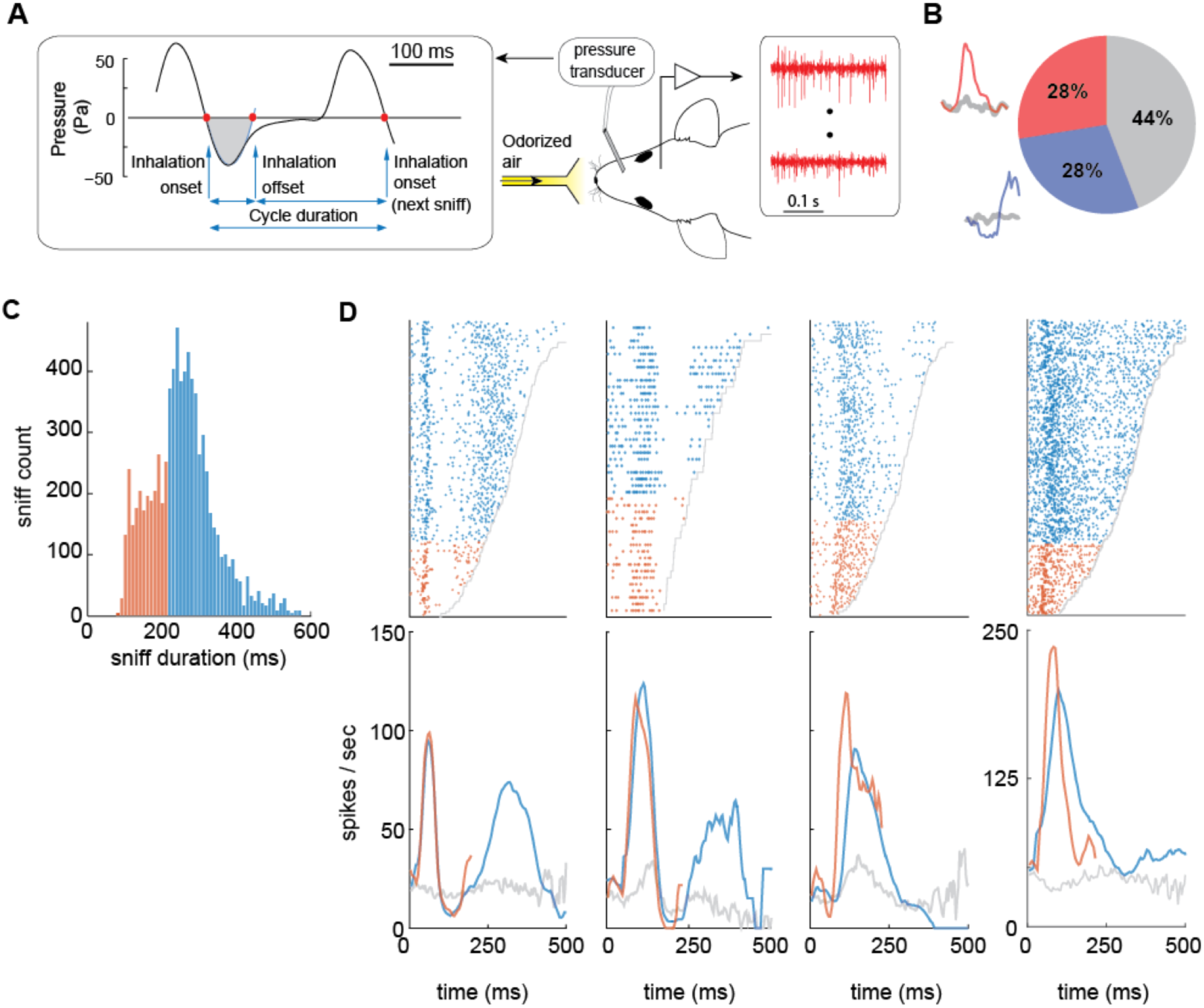
Differences in MT cell responses between fast and slow sniff cycles. **A.** Schematic of the experiment. A head-fixed mouse implanted with intranasal cannula and a multi-electrode chamber was positioned in front of the odor delivery port. Left, pressure waveform of a typical breathing cycle. Red dots indicate the inhalation onsets and offsets. The blue line is the parabolic fit to the first minimum after the inhalation onset. The sniff offset was defined as the second zero crossing of the parabolic fit. The gray shaded area marks the inhalation interval. **B.** Distribution of different response types observed in the data. Next to the pie chart are examples of initially excitatory (red) and initially inhibitory (blue) responses. Gray lines represent spontaneous activity. **C.** Sniff duration histogram for all mice averaged across multiple sessions for the 1^st^ odorized sniff cycle. Sniffs are split into two groups: fast (< 221 ms; orange) and slow (> 221 ms; blue). **D.** Raster (top panels) and PSTH (bottom panels) plots of the first odorized sniff cycle for four MT cells in response to an odor stimulus. Trials in the raster plots are ordered bottom to top by the increase in the duration of sniff cycle. Gray line shows end of the sniff cycle. Raster plots and PSTHs for fast and slow sniffs are marked by orange and blue colors, respectively. Gray PSTHs are average spontaneous activity of MT cells during all unodorized sniffs.

To visualize MT cell odor response dependence on sniff duration, we aligned responses by the beginning of the first inhalation after the onset of odor presentation. Trials in the raster plots were ordered by the duration of the first odorized sniff cycle. These raster plots reveal the clear dependence of MT cell responses on sniff frequency (Fig. 2D). While earlier temporal features of MT response are preserved across different sniff durations, the later features are often absent on trials with short sniff cycles (Fig. 2D). Thus, the temporal pattern across the full sniff cycle depends on the sniff duration.

## Amplitude and latency of the response but not spike count depend on sniff duration

Visual inspection of the raster plots (Fig. 2D) suggests that responses during the early part of the sniff cycle are more similar for fast and slow sniffs than the later part. To quantify this observation, we estimated the distribution of sniff durations for all mice and all sessions and divided the sniffs into two groups, fast sniff trials with the sniff duration in the left third of the distribution, and slow - in the right two thirds of the distribution (Fig. 2C). The boundary between these groups was 221 ms. (see Methods, Fig. 2C). MT cell activity during short and long sniff cycles was averaged separately (Fig. 2D). MT cell responses during slow sniff cycles contain on average 2.3 times more spikes than responses during fast sniff cycles. However, the average spike count during the first 221 ms remains almost the same (Fig.3A). The distribution of the spike count differences for fast and slow sniffs for all cell-odor pairs is centered around zero (Fig. 3A, insert, p = 0.43, Wilcoxon signed rank test). Thus, spike count over this early time window is invariant to changes in sniff frequency.

**Figure 3.**
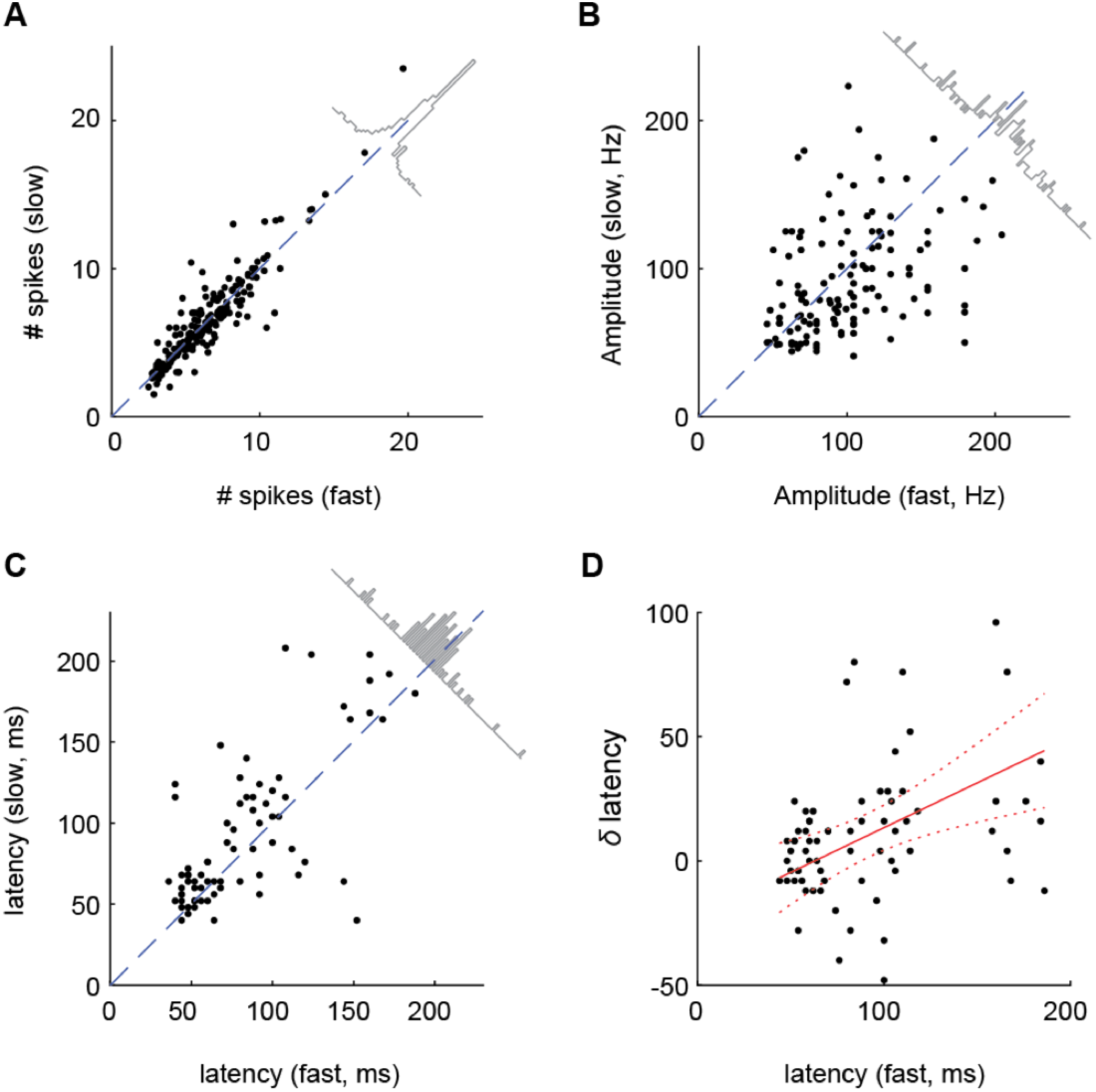
Comparison of response features for fast and slow sniff durations. **A.B.C.** Comparison scatter plots of average spike count during the first 221 ms (A), average peak amplitude (B), and average response latency (C) for fast vs slow sniff cycles. Each dot represents comparison for a single cell-odor pair. Diagonal plots are distributions of differences of corresponding values. Spike counts are statistically indistinguishable for slow and fast sniffs. On average, peak amplitudes are larger (p = 0.013; Wilcoxon signed rank test) and latencies are shorter (p = 0.007; Wilcoxon signed rank test)) for fast sniffs compared to slow sniffs. **D.** Shift in latency of responses between slow and fast sniffs as function of response latency. Red solid line is a linear regression fit to the data. Red dashed lines mark the limits of the 95% confidence interval (slope = 0.35).

Integration of spiking activity over a long time widow (∼200 ms) may not serve as an informative cue for many odor driven behavioral decisions, which are often relatively fast (Resulaj and Rinberg, 2015; Wilson et al., 2017). Thus, we looked into finer features of MT cell responses, such as the amplitude of the response, which is a combined characteristic of instantaneous firing rate and spike timing precision, and the latency of the response (Fig. 2D). We calculated these parameters for fast and slow sniffs separately. We present the data for individual cell-odor pairs as scatter plots of peak amplitudes (Fig. 3C) and peak latencies (Fig. 3D) for fast and slow sniffs. The response amplitude is slightly lower for slow sniffs than that for fast sniffs (Fig.3B, insert, *δ*A=5.7 Hz, p = 0.013, Wilcoxon signed rank test). The response latencies also differ between fast and slow sniffs – fast sniff latencies are slightly shorter that those of slow sniffs (Fig. 3C, insert, *δ*τ =10.4 ms, p = 0.007, Wilcoxon signed rank test). Further analysis of the response latencies showed that latency differences *δ*τ between slow and fast sniffs increases with latency (Fig. 3D, regression parameters: slope=0.36+/-0.11, intercept=-22.6+/-11.2). Thus, fine temporal features of the odor response vary with the sniff duration.

How significant are these sniff duration-dependent differences in response latency and peak amplitude? To answer this question, we compare the time and amplitude changes due to sniff variability to those due to concentration changes. Based on our previous work (Sirotin et al., 2015), a time difference of *δ*τ≈10 ms corresponds to a 3-fold concentration change, and an amplitude difference of *δ*A≈6Hz corresponds to a 2- fold concentration change. So, the temporal features of MT cell responses vary with sniff duration as much as they vary for 2-3 fold concentration differences. These observations raise two questions. First, what is the mechanism responsible for the sniff waveform dependent variability? And second, is there a representation of the odor response which is invariant to the sniff duration? These two questions are related: defining the mechanism will allow us to find a better representation, and vice versa: finding a better representation will help reveal the mechanism.

Next, we formulate a simple model based on the fluid dynamics of odor propagation in the nasal cavity which may explain the sniff dependent variability. We then compare our model with the previously proposed representations described above. For quantitative comparison, we propose a metric of how well these representations reduce sniff waveform related odor response variability. Lastly, we discuss the applicability of such transformations for analysis of olfactory coding.

## Fluid dynamic model

We observed that faster sniffs evoke earlier odor responses with higher amplitude. Also, on average, faster sniffs have larger pressure amplitude changes (Fig. S2; Walker et al., 1997; Youngentob et al., 1987). We assume that the pressure measured via cannula is a good proxy of the air velocity inside the nasal cavity. Given this assumption, we propose a fluid dynamics-based model which can explain the dependence of response latency on a sniff waveform.

Let’s assume that response latency, τ, of a MT cell odor response consists of two terms: first is a fluid dynamic term that depends on the sniff waveform, τ_*f*_[*u*(*t*)] (where, *u*(*t*) is the air velocity waveform, and square brackets symbolize the dependence on the whole waveform function, rather than instantaneous values). Second term, τ_*n*_, is the delay due to neural processing which may depend on the odor and a specific MT cell, but is independent of the sniff waveform:

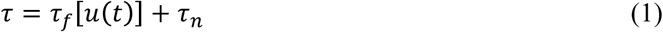

To model the dependence of τ_*f*_ on the sniff waveform, we propose that odor is carried into the nose by an air flux with time-dependent velocity *u*(*t*) (Fig. 4A). Suppose, the receptors are located at an approximate distance *L* from the nasal opening. Then the relationship between air velocity *u*(*t*) and distance *L* is following:

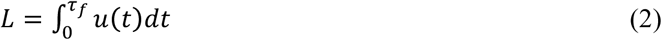

**Figure 4.**
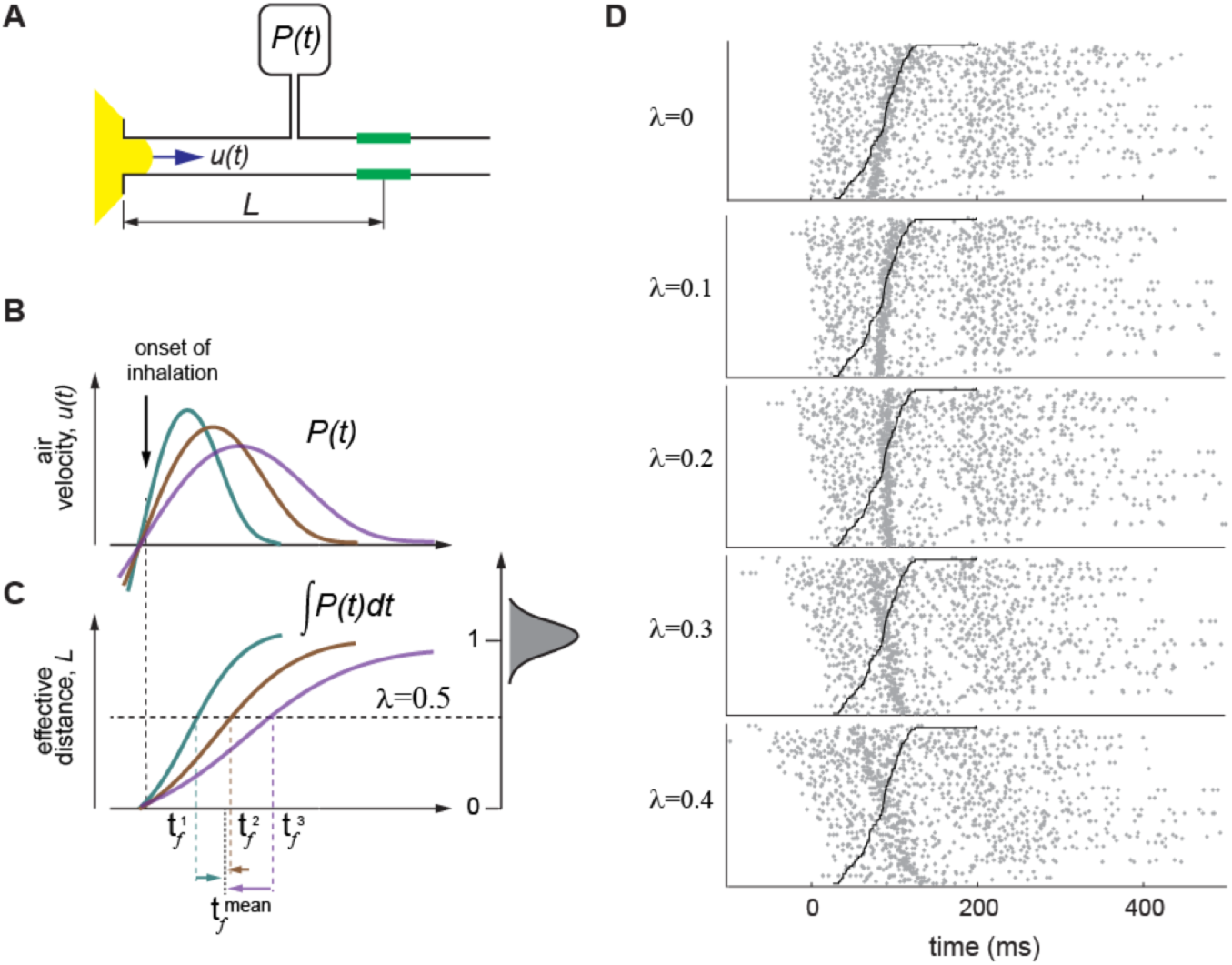
A fluid dynamics model explaining latency of MT cell activation **A.** Simple physical model of odor molecules delivery to the receptors. During inhalation, odorized air (yellow) is drawn into the nose with velocity *u(t)*. The OSNs (green) are located at an average distance *L* from nasal opening. Air velocity is proportional to pressure measured via cannula *u(t) ∼ P(t)*. **B.** A schematic of pressure signal temporal profiles, *P(t)* for three representative sniff cycles. **C.** Integrals of pressure signals: 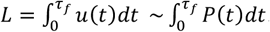. Left panel: a schematic distribution of values of integrals over the whole inhalation interval normalized to their mean value, *λ*=*L*/*L*_*max*_>. For a given value of an integral, for example, λ=0.5 in normalized coordinates, we estimate time points 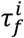for which: 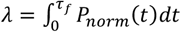, here *P*_*norm*_(*t*) are normalized pressure traces. **D.** Raster plots of spiking data from a representative cell, aligned to different values of λ. Black line shows end of the inhalation.

In our model, *L* does not depend on the sniff waveform, and the velocity is proportional to the measured pressure: *u*(*t*)∼*P*(*t*). Under these conditions, during faster sniff cycles, air velocity is higher, therefore it takes less time τ_*f*_ to odor molecules to pass the same distance *L*. While *L* is unknown we can treat it as a model parameter which is estimated by minimizing neural response variability. The detailed procedure is shown in Figure 4B, C and in Methods. Briefly, for a given fraction, *λ*, of normalized length, (*λ α L*), and for each sniff *i*, we find the fluid dynamic time, 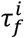. Then we estimate an average fluid dynamic delay 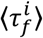 across all trials, and shift spiking activity during trial *i* by a time interval 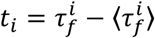. We repeat this procedure for different *λs* between 0 and 1. Then, using log-likelihood estimation, we find the value of *λ* that minimizes trial-by-trial variability of the neural response. The transformation of spike trains for different values of *λ* is shown in Fig. 4D for one cell-odor pair (also see Fig. S3). On average, the best transformation of odor responses is achieved for *λ*=0.3 (Fig. S4).

This model-based procedure reduces odor response variability compared to an original representation in time relative to the onset of inhalation, which corresponds to *λ*=0. In sensory systems, neuronal responses are often tightly locked to the time of stimulus presentation (Richmond et al., 1990). In olfaction, this time may be the moment when odor concentration crosses a receptor’s threshold. This timing varies with the sniff pattern. Although our model is oversimplified, it may allow us to remove the influence of sniffing on spike pattern and better understand sniff invariant odor processing. Next, we perform model based comparison of different representations to find a transformation that best describes sniff invariant odor representation.

## Model comparison

To infer the best sniff invariant odor representation, we apply different models to transform individual spike trains based on the corresponding waveforms, and estimate an average odor response. The optimal transformation should minimize trial to trial response variability. Using maximum likelihood estimation (MLE), we evaluate which of the average responses better fits experimental data (see Methods). We consider the following spike time transformations, most of which were previously discussed in the literature, while the last one is proposed here (Fig. 5A):

**Figure 5.**
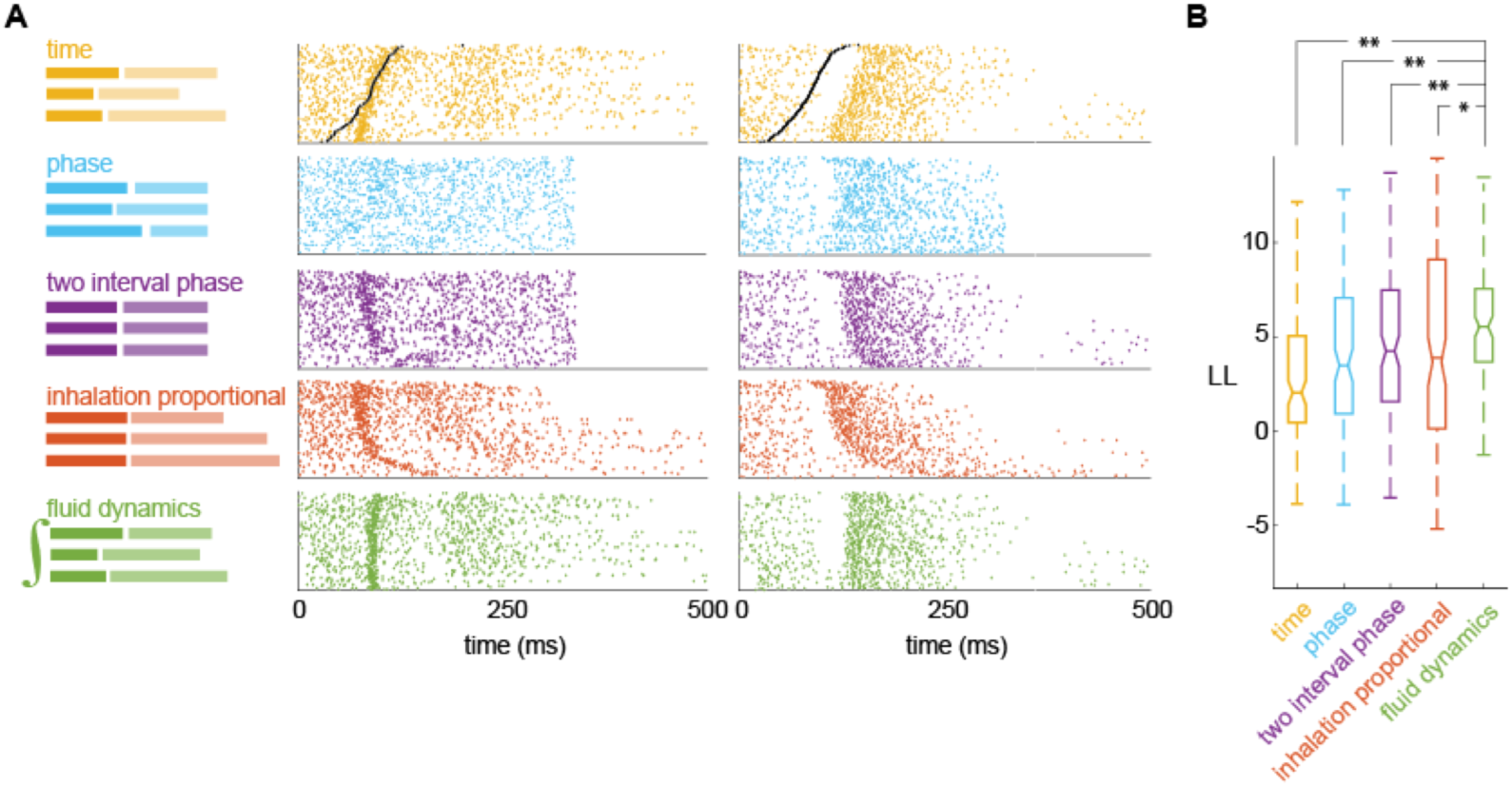
Model comparison of different alignment methods Left: Schematics of the effect of each of the transformation on sniff cycles. Right: raster plots of responses of two cell-odor pairs aligned according to each schematic. **B.** Log-likelihood of each model across the population of responses. Pairwise comparisons were done using the Wilcoxon signed rank test (FD vs. time comparison, p=5.7e-19; FD vs. phase comparison, p=9.5e-04; FD vs. two interval phase comparison, p=9.9e-03; FD vs. inhalation proportional comparison, p=3.5e-02).

1. *Time model*. Spike times aligned to onset of inhalation, with no further transformation.
2. *Phase model*. Spike times relative to onset of inhalation are scaled proportionally to the duration of the whole sniff cycle.
3. *Two-interval phase model.* The sniff cycle is split into the inhalation interval and the rest of the cycle. Spike times during the inhalation interval are scaled proportionally to the duration of the inhalation, while spikes times occurring in the rest of the sniff cycle are scaled proportionally to the duration of that interval (Shusterman et al., 2011).
4. *Inhalation proportional model.* Spike times during the whole sniff cycle are scaled proportionally to the duration of the inhalation (Arneodo et al., 2017).
5. *Fluid dynamics based model.* Spike times are shifted by a specific value determined by the sniff waveform integral in that trial (see Fig. 4). Note that only this model has an additional adjustable parameter, optimization of this parameter was done using the same MLE procedure.

An example of transformation for all 5 models of one cell-odor response is shown in Fig. 5A. To fit each model, we transform the spike trains in individual trials according to that model’s alignment rule (Fig. 5A) and calculate PSTHs using the transformed spike trains. To evaluate the goodness of fit of different models, we evaluate their average log-likelihood in held out sniff cycles. To this end, we first inversely transform each model’s PSTH according to the duration parameters of a given sniff cycle using interpolation. We then evaluate the Poisson process log-likelihood, given the inversely transformed PSTH, for the spike train during that sniff cycle (Fig. 5B). Finally, we average the calculated single-sniff log-likelihoods for a model over all sniffs to obtain that model’s average log-likelihood. We performed this analysis on 192 excitatory responses each consisting of at least 30 trials per response. On average, the FD model better predicted the time course of instantaneous firing rates than any of the other models (Fig. 5B, see Methods for details). Pairwise comparisons were done using the Wilcoxon signed rank test on population log-likelihoods (FD vs. time comparison, p=5.7e-19; FD vs. phase comparison, p=9.5e-04; FD vs. two interval phase comparison, p=9.9e-03; FD vs. inhalation proportional comparison, p=3.5e-02). However, we need to note that the FD model is the only model which has a fitting parameter, which potentially makes a better fit. Among no-fitting parameters models the best is two-interval phase model, which we used previously in our studies to discover higher temporal precision of odor responses (Shusterman et al., 2011). The difference between two-interval phase and inhalation proportional models is insignificant (*p=0.12)*, and may be different for different data sets (Arneodo et al., 2017).

## Decoding of odor concentration in FD transformation is better than in real time

If a model based transformation of spike trains reduces sniff waveform related variability, then stimulus decoding using that transformation should be improved. Response profiles of the same cell across odorants are highly diverse (Shusterman et al., 2011). Therefore, odor identity can be decoded with high success rate irrespective of how or whether they are transformed (Bathellier et al., 2008; Cury and Uchida, 2010; Shusterman et al., 2011). In contrast, the type of transformation should play a more significant role in decoding odor concentration, which is encoded mainly by the fine temporal structure of spike trains (Bolding and Franks, 2017; Cang and Isaacson, 2003; Sirotin et al., 2015). As we pointed out above, sniff waveform-related variability is similar to a 2 to 3-fold uncertainty in concentration estimation. To test if the FD alignment allows for better decoding of odor concentration than time alignment, we evaluated the accuracy with which responses of MT cells to two odor concentrations (Fig. 6A, B) can be discriminated in each alignment, on a trial-by-trial basis (See Methods). We performed this analysis on 93 cell-odor-concentration triplets. On average, discrimination success rate between odor concentrations is better after FD transformation than in time coordinates: 70.2% vs 78.3% (t-test, p = 8e-03; Fig. 6C).

**Figure 6.**
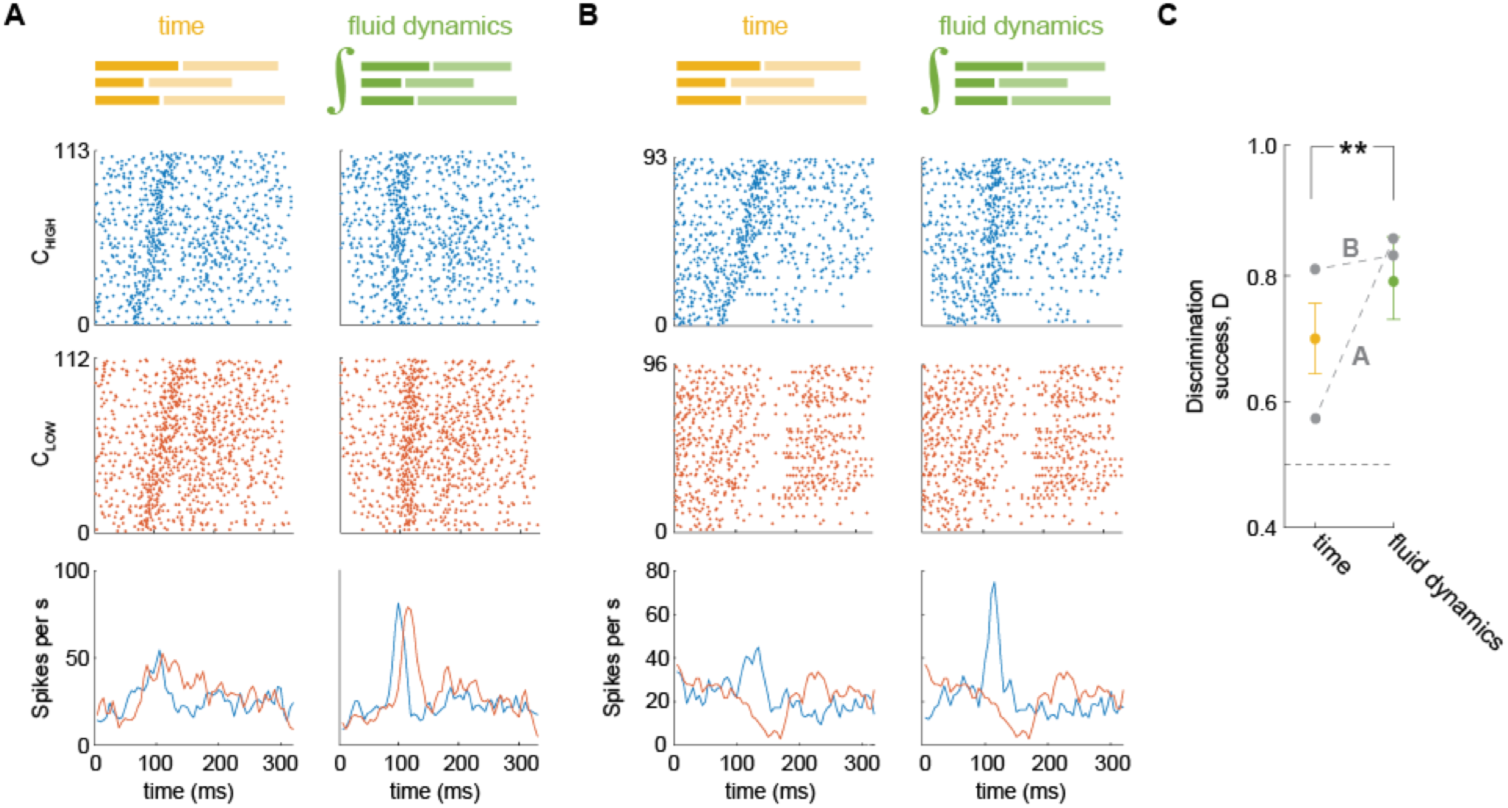
Discrimination among odor concentrations by MT cells in time vs. FD transformation. A. B. Raster and PSTH plots of MT cell responses to high (blue) and low (red) odor concentrations in time and in FD transformation. **C.** Concentration discrimination performance across the population in time (yellow) and FD (green) alignments. Gray dots and lines show discrimination performance for odor concentration pairs from (A) and (B). The horizontal dashed line is chance level performance.

## Discussion

Sensory systems must extract accurate representations of the outside world despite variability in sampling behavior. We studied whether odor representation is invariant to changes in sampling frequency. We found that behavioral performance in an odor concentration discrimination task does not depend on sniff frequency, suggesting that odor representations are indeed sniff frequency invariant. To identify neural signatures of this invariance, we studied responses of MT cells to different odors and concentrations. The amplitude and the latency of MT cells’ responses varied systematically with sniff duration. We proposed a FD model that explains this response variability and provides the best description of the odor responses. Further, transforming odor responses based on the FD model makes the responses invariant to sniff waveform variability, and allows much more accurate concentration discrimination.

## Limitations of proposed fluid dynamic based model

We propose a very simple fluid dynamics model for odor propagation in the nose. Obviously, it does not capture all the complexity of this process, however, it does provide a reasonable explanation for how variability in sniff waveforms contribute to variability in MT cell odor responses. A few questions remain to be discussed: What are the limitations of our simple model? Is it compatible with our results on the concentration dependence of the odor responses? Lastly, given a proposed model of odor delivery to the receptors, how does coding of odor identity and concentration work?

According to our FD model, the front of odor-laden air propagates in the nose with a velocity which increases and then decreases during an inhalation, approximately proportional to nasal cavity pressure (Fig. 7A). In a simple version of this model, the propagating front has a sharp boundary, such that odor concentration at the receptors’ location will rise as a step function (Fig. 7B). In reality, the front boundary is not sharp (Nachbar and Morton, 1981). Rather, it spreads as the front propagates along the nasal cavity (Fig. 7C). There are few factors which contribute to this effect: 1) Due to friction or air viscosity, flow close to the edge of the channel moves more slowly than flow near the center of the channel; 2) Due to adsorption by respiratory and olfactory epithelium, odorant concentration decreases along the way. Adsorption depends strongly on the physical-chemical properties of an odorant, and may significantly differ for hydrophobic and hydrophilic molecules. Thus, concentration increases gradually in any given location of the epithelium (Fig. 7C). In addition, receptors of the same type are not localized to one point, but rather are spread through the epithelium, which slows the effective concentration rise even further. In future work, these effects can be modeled or simulated. At present, our FD model provides a simple explanation of how odor propagation in the nose causes MT cell responses to vary with sniff waveform.

**Figure 7.**
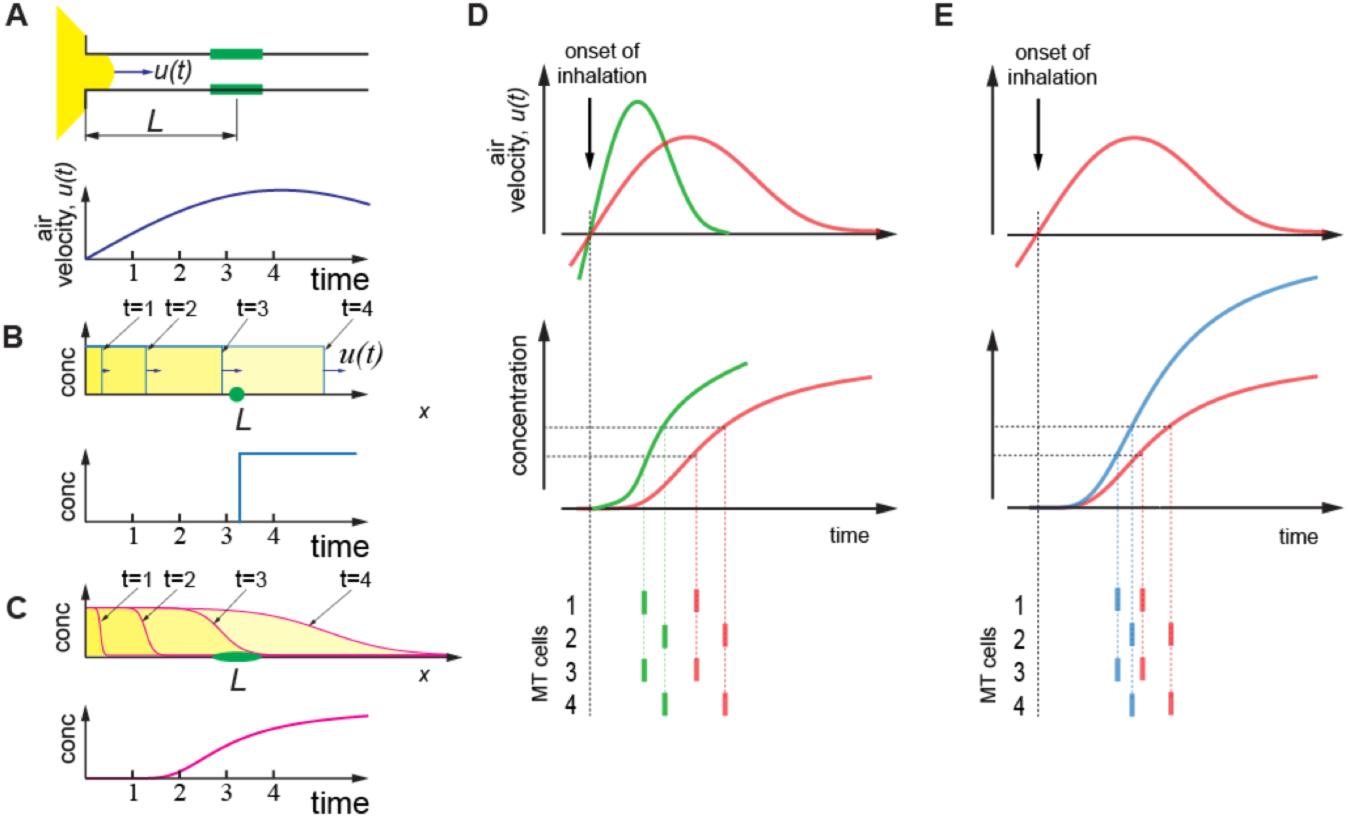
Effect of sniff rate and odor concentration on spiking latency **A.** Top: Schematics of the fluid dynamics model. Same as in Fig. 4A. Bottom: schematic of the air velocity temporal profile in the nose during inhalation. **B.** Schematics of the odor concentration wave-fronts at different times from the beginning of inhalation in a simple model presented at **A** (top) and a correspondent temporal concentration profile at a specific point in the nasal cavity positioned at the distance *L* from the nasal cavity opening (bottom). **C.** More realistic case of odor molecules’ delivery to the receptors, in which friction and epithelial adsorption smear the odorized air front. Top: Propagation of an odorized air front inside the nasal cavity according to the model from (A). Bottom: rise time of odor concentration at the receptor location. **D.** Effects of increase in sniff frequency on spiking latency of MT cells. Top: air velocity profile during fast (green) and slow (red) inhalation. Center: Corresponding temporal profile of odor concentration at the receptor location. Bottom: MT cell spiking driven by corresponding OSNs which are activated at different concentration levels. **E.** Same as (D) for two different concentrations of the same odor.

The next important step in our model is to define how OSN signals are triggered and conveyed to the OB. Most simply, we may assume that OSNs spike when the concentration of a ligand reaches some threshold level. Under this assumption, in the case of sharp boundary waveform propagation (Fig. 7B), no matter what the external odorant concentration is, the ligands will arrive at the receptor location at the same time. Therefore, time of activation of OSNs and downstream MT cells will be concentration independent. However, if ligand concentration at the epithelium gradually increases (Fig 7C), higher stimulus concentrations will activate OSNs earlier relative to the sniff cycle (Fig. 7E) which is consistent with our previous observations (Sirotin et al., 2015). Thus, the qualitative extension of FD model predicts that the latency of OSN activations and following MT cell responses will be shorter for higher stimulus concentration, which will manifest in a smaller effective distance, *λ*. Based on our measurements of the responses of the same cell to the same odor with different concentrations, at higher concentration *λ* is smaller than that for low concentration (Fig. S6).

## Coding of odor identity and odor concentration

Given the FD model, how might odor coding work? MT cells in the OB are the first recipients of sensory information from the OSNs and the only cells that project their axons outside the bulb to olfactory cortices (OCs). The information that OCs receive about odor stimulus is a sequence of MT cell activations (Hopfield, 1995; Margrie and Schaefer, 2003; Schaefer and Margrie, 2007; Shusterman et al., 2011; Wilson et al., 2017). This code can be read based on synchrony between multiple simultaneously activated MT cells (Bolding and Franks, 2017; Franks and Isaacson, 2006; Haddad et al., 2013; Sanders et al., 2014). During a given 10 ms time window in the sniffing cycle, hundreds of MT cells can be activated simultaneously in response to an odor (Shusterman et al., 2011). The anatomical finding that MT cells from multiple glomeruli converge to one pyramidal cell in the cortex (Miyamichi et al., 2011), and electrophysiological recordings demonstrating that pyramidal cells integration time window is ∼10 ms (Davison and Ehlers, 2011; Poo and Isaacson, 2009), as well as work in anesthetized animals on sensitivity of cortical neurons to timing of MT cells activation (Haddad et al., 2013), are consistent with a synchrony-based model of reading the MT cells’ code.

The recently-proposed primacy coding hypothesis (Bolding and Franks, 2017; Wilson et al., 2017), states that identity-carrying responses occur during an early and narrow time window after inhalation onset. This coding scheme is robust to sniff waveform variability, because the earliest-activated OSNs, and following MT cell responses would remain the same for different sniff durations and are subjects to less variability (Figs. 2 and 3).

How does odor concentration encoded? Both sniff waveform and odor concentration affect OSN activation latencies. The sniff waveform regulates the speed of odor propagation, which affects mostly the onset, and also the steepness of the concentration profile (Fig. 7D). Concentration changes will only affect the steepness (Fig. 7E). An earlier or steeper rise of odor concentration, whatever its cause, will compress the timing of odor responses, as well as shift them earlier in the sniff cycle. Nevertheless, concentration judgments maintain constancy despite varying sniff parameters (Fig 1); (R. Teghtsoonian and M. Teghtsoonian, 1984). How might the olfactory system achieve this perceptual constancy? One possibility is that the olfactory system has access to the sniff waveform parameters. This idea is supported by the finding that mice can report the latency of OSN activation relative to inhalation (Smear et al., 2011). To generate a sniff representation, the mouse may use reafference, as many OSNs are mechanosensitive (Grosmaitre et al., 2007), and could therefore sense airflow in the nose. Another possible mechanism for representation of the sniff waveform is through efference copy, although there is no evidence to support this claim, nor is there a candidate anatomical pathway between breathing centers and the olfactory system, to our knowledge. However, it also remains possible that the animal does not use timing information. Higher odor concentration leads to larger number of activated OSNs (Rubin and Katz, 1999; Stewart et al., 1979).

There are multiple normalization mechanisms in the OB, though it does not preclude existence of channels transmitting overall level of activation. However, the latter mechanism probably requires accumulating signals during longer time period, which is not well supported by behavioral data (Resulaj and Rinberg, 2015; Wesson et al., 2008; Wilson et al., 2017). Another possibility for concentration encoding may arise from the fact that based on our arguments beyond a simple FD model, the concentration increase may evoke a stronger temporal compression of the spike trains, then that evoked by faster sniffing (Fig. 7E). This will in turn increase the level of synchrony, which has been observed in cortical recordings (Bolding and Franks, 2017). Quite possibly all of the above mechanisms participate in multiplexed transmission of the information about concentration. Future work will be required to dissect and define their contributions.

## Author contributions

R.S. and D.R. conceptualized the study; Y.B.S. performed behavioral experiments; R.S. and Y.A. analyzed the data; R.S. and D.R. interpreted the data; R.S., M.S., and D.R. wrote the manuscript.

## Acknowledgments

We would like to acknowledge Chris Wilson and Avinash Bala for discussions, Joel Mainland and Kai Zhao for comments on the manuscript. This work was supported by the Israel Science Foundation grants 816/14 and 2212/14 (R.S), Marie Curie Career Integration grant 334341 (R.S.) and by NIH/NICDC grants: R01DC014366 and R01DC013797 (D.R.).

## Methods

### Behavioral experiments

All experimental procedures were conducted according to the US National Institutes of Health Guidelines for the Care and Use of Animals. Behavioral experiments were performed at Rockefeller University and were reviewed and approved by the Institutional Animal Care and Use Committee. A total of 4 rats were used in this study. Animals were housed in a 12 h light/dark cycle and behavioral testing was performed during the dark cycle.

<u>Odors and Odor Delivery.</u> For behavioral experiments, we used the following odors pentyl acetate, ethyl acetate, (R)- (+)-limonene, (+)-α-pinene, phenyl ethyl alcohol, and decanal. Odors were selected based on rapid delivery kinetics from olfactometer (Martelli et al., 2013). For behavioral studies we used the same odor delivery setup and odor delivery principles as for electrophysiological experiments described above.

<u>Behavioral Apparatus.</u> Behavioral experiments were conducted in a custom made behavior box (Bodyak and Slotnick, 1999). The behavior box was built from acrylic and stainless steel to avoid odor contamination. Air within the box was exchanged by a fan at a rate of 15 L/s. We used a three ‘ports’ design (Uchida and Mainen, 2003). The central port was used for odor delivery and two side ports were set up to provide water reward (0.023 ml). The central port was a custom made Teflon odor port, connected with the output of the olfactometer. To ensure consistent nose placement in the odor stream across trials, the orifice of the odor port was narrow (8 mm). To reduce outside noise and light, the box was surrounded by an additional aluminum enclosure with foam padding. We used infra-red camera to monitor animals externally.

<u>Initial training.</u> We used Long Evans rats from Charles River Laboratories. Training began when rats reached a weight of at least 200 g. Throughout training, rats were water restricted. At least once a week animals had *ad libitum* water access. After each behavioral session, rats were given 15 minutes of *ad libitum* water access to ensure adequate hydration. Weight and water intake were monitored daily to ensure normal growth rates.

Initially, rats learned to poke into the reward ports positioned on either side of a central odor port to obtain a water reward (1 session). Then, animals were trained to initiate trials by nose poke (minimum poke time: 50 ms) into the central odor port prior to getting a reward at either of the reward ports (1-4 sessions). After this behavior reached stable performance, rats were shifted to an odor/no odor task: an odor was introduced randomly on some trials. Reward was delivered to the left water port for odor present trials and to the right port for no odor trials. No effort was made to correct for any biases or alternation strategies. Rats readily learned to ignore the unrewarded port on each trial (1-2 sessions) after which point reward was no longer given for ‘incorrect’ pokes (1-2 sessions). At this point, rats were trained to perform the concentration classification task. Rats typically reached asymptotic performance in ∼3 sessions. Collection of experimental data started after the rats established stable psychometric functions: between 10 to 20 days after initiation of training.

<u>Sniffing cannula and pressure sensor implantation.</u>

Rats were implanted with sniffing cannula in the same procedure to sniffing cannula implantation in mice, described below. A pressure sensor (24PCEFJ6G, Honeywell) was mounted on the animals head. It was attached to the scull with dental cement. The cannula was connected to pressure sensor via short polyethylene tubing (801000, A-M systems). After surgery, a rat was given at least 7 days for recovery.

<u>Concentration classification task</u>

Rats were run daily during the weekdays on the concentration classification task. On different days, rats were tested with different odors at different dilutions (mineral oil) in the vial. After switching between odor stimuli, rats took 1 to 4 sessions to reach criterion performance (perceived category bound less than 0.1 log units away from the trained category bound). To ensure enough trials per concentration, for this analysis we only used session where the rats performed over 700 trials.

In each session, rats initiated trials by poking their nose into the odor port triggering odor delivery from the olfactometer for sampling. From the onset of the poke, it took 50 ms for the odor to reach the nose. Delivered odor concentration was varied by flow dilution in eight steps across one order of magnitude. On each trial, rats were free to sample the odor for a maximum of one second. Later for analysis we excluded trials in which sampling was longer than one sniff cycle. Rats were rewarded for going to the left reward port for the four ‘high’ concentrations and the right port for the four ‘low’ concentrations. Trials on which the animals did not stay in the odor port at least 50 ms or didn’t access the reward port were not rewarded or analyzed further. Failures to seek reward made up less than 1% of all trials. For each trial, behavioral choice (left or right), odor sampling time, sniffing pressure and response time were recorded. Odor sampling time was counted as the duration of time the rat spent in the odor port. After each trial, a three second inter-trial-interval was enforced.

### Electrophysiological experiments

<u>Experiments</u>. The analysis presented in this manuscript was performed with electrophysiological data obtained in the experiments described in previous publications (Shusterman et al., 2011; Sirotin et al., 2015).

<u>Animals</u>. Electrophysiological data were collected in ten C57BL/6J mice and three OMP-ChannelRhodopsin-YFP heterozygous mice that have a targeted insertion in the OMP locus (Smear et al., 2011). All mice had at least one normal copy of OMP. No differences were observed between these two groups of mice. Subjects were 6–8 weeks old at the beginning of behavioral training and were maintained on a 12 h light/dark cycle (lights on at 8:00 P.M.) in isolated cages in a temperature- and humidity-controlled animal facility. All animal care and electrophysiological procedures were in strict accordance with a protocol approved by the Howard Hughes Medical Institute Institutional Animal Care and Use Committee.

<u>Electrophysiology</u>. Mitral/tufted cell spiking activity was recorded using 16 or 32 channel Silicon probes (NeuroNexus, model: a2x2-tet-3mm-150-150-312 (F16), a4x8-5mm 150-200-312 (F32)) or using a home-made microdrive with 16 individually adjustable 3 MOhm PtIr electrodes (MicroProbes, MD, USA). Cells were recorded from both ventral and dorsal mitral cell layers. MT cells were identified based on criteria established previously (Rinberg et al., 2006). The data was acquired using a 32 channel Cheetah Digital Lynx data acquisition system (Neuralynx, Tucson, AZ) with opened band-pass filters (0.1 - 9000 Hz) at 32.556 kHz sampling frequency.

<u>Sniff monitoring</u>. Sniff signals were monitored via a thin, stainless cannula (Small Parts capillary tubing, Ga 22-23) implanted in the nasal cavity. In between experimental recordings, the cannula was capped with a dummy plug. We used polyethylene tubing (801000, A-M systems) to connect the cannula to a pressure sensor (MPX5050, Freescale Semiconductor or 24PCEFJ6G, Honeywell). The signal from the pressure sensor was amplified with a homemade preamplifier circuit and was recorded in parallel with electrophysiological data on one of the analog data acquisition channels.

<u>Surgery.</u> Mice were anesthetized and implanted with a head fixation bar, pressure cannula, and Microdrive with mounted Silicon probe. For the sniffing cannula implantation, a small hole was drilled in the nasal bone, into which the cannula was inserted and fastened with glue and stabilized with dental cement. For the electrode chamber implantation, a small craniotomy (1mm2) was drilled above the olfactory bulb (contralateral to the sniffing cannula position) and dura mater was removed. The Silicon probe was inserted, and then the electrode chamber was fixed by dental cement to the skull, posterior to the olfactory bulb. A reference electrode (76 um PtIr PFA-insulated wire, A-M Systems, WA USA) was implanted in cerebellum. A mouse was given at least 5 days after a surgery for recovery.

<u>Odor delivery.</u> We used multiple odorants obtained from Sigma-Aldrich (St. Louis, MO). The odorants were stored in liquid phase (diluted 1:5 in mineral oil) in dark vials. The odorant concentration delivered to the animal was additionally reduced tenfold by air dilution. Odorants used in the study are: acetophenone, amyl acetate, behzaldehyde, butyric acid, decanol, ethyl acetate, ethyl tiglate, 1-hexanol, hexanoic acid, hexanal, 2-hexanone, hexyl acetate, R-limonene, isopropyl tiglate, methyl benzoate, methyl salicylate, 1- octanol, 2-undecanone.

For stimulus delivery we used a 9 odor air dilution olfactometer. To prepare different concentrations, air flow through the selected odor vial was mixed with clean air in a variable ratio configured electronically on each trial by mass flow controllers (Bronkhorst or Alicat Scientific). The air flow through each odor vial could vary between 10 ml/min to 100 ml/min. This odorized air was mixed with a 990 ml/min to 900 ml/min clean air carrier stream, so that total flow coming out of olfactometer is always 1L/min. A steady stream of 1 L/min of clean air was flowing to the odor port all the time except during stimulus delivery. After an odor stimulus was prepared, a final valve (4 way Teflon valve, NResearch) delivered the odor flow to the odor port, and diverted the clean airflow to the exhaust. All flows and line impedances were tuned to minimize the pressure shock due to line switching and minimize the time of odor concentration stabilization after opening the final valve. The temporal odor concentration profile was checked by mini PID (Aurora Scientific). The concentration reached a steady state 25-40 ms after final valve opening.

<u>Stimulation protocol.</u> After recovery from surgery, the animal was placed in the head fixation setup. First 2 short sessions were used to adapt the animals to head fixation in the setup. After habituation, we began recording sessions. In each trial, one out of 2-10 odors was delivered in pseudo-random sequence with an average inter-stimulus interval of 7 sec. Odor delivery was triggered by the offset of inhalation. Since odor can’t enter the nose during exhalation phase, the duration of exhalation interval allowed enough time for the odor stimulus to reach a steady state of concentration by the time the animal begins next inhalation. Each session usually contained 300 - 600 trails and lasted for 45-90 min.

**Data analysis**

All analysis was done in Matlab (Mathworks Natick, MA).

<u>Psychometric functions.</u> Psychometric curves were fitted to behavioral data using maximum likelihood (Wichmann 2001). The fitted psychometric function *ψ* had the following form:

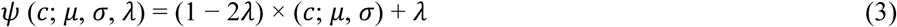

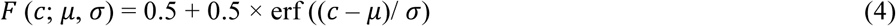

where *c* is the odor concentration, *μ* is the categorical boundary, *σ* is the noise, and *λ* is the guess rate. Goodness of fit was assessed using deviance, computed as the log likelihood ratio between the fitted and the saturated models (Wichmann and Hill, 2001a). Confidence intervals of the parameters were estimated by the bootstrap method based on 250 repetitions (Wichmann and Hill, 2001b).

<u>Spike extraction.</u> Acquired electrophysiological data were filtered and spike sorted. For spike sorting we used the M-Clust program (MClust-3.5, A. D. Redish et al) and a software package written by Alex Koulakov.

<u>Odor responses.</u> We compared the distributions of the neuronal activity with and without odors. Distribution without stimulus was sampled from 5 sniffs preceding the odor delivery across trials (approximately 1500- 2000 sniffs for each session). For all neurons, we determined the distribution of spikes for each stimulus on the first sniff after stimulus onset. To establish if a cell responded to particular odorant, we tested whether the cumulative distribution of spike counts differed significantly between background and stimulus distributions (Kolmogorov-Smirnov test, p-value 0.05). The cumulative distributions were calculated using spike times binned at 1 ms. The response was considered excitatory/inhibitory if the first deviation from the background distribution was positive/negative.

<u>Fitting FD model.</u> To apply fluid dynamics model to MT cells responses, for each trial *i* we align all spike trains to a specific time moments during inhalation interval, 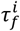, defined by the condition (Fig. 4A):

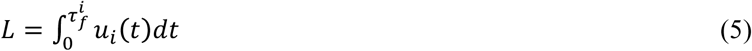

In this equation, *L* is an effective distance between a nostril opening and the location of OSNs, activation of which lead to a given MT cell response. *L* is unknown, but we assume it to be identical for all sniffs for a given cell-odor trials. *u*_*i*_(*t*) is an air velocity at trial *i*, which we assume to be proportional to a pressure signal measured in the experiment: *u*_*i*_(*t*)=𝛼*P*_*i*_(*t*). During faster sniff cycles, it takes less time τ_*f*_ for odor molecules to pass the same distance *L*.

Under these assumptions, we are searching for *L,* which minimizes the neural response variability. As we do not know the value of the coefficient α, we perform a search in normalized distance variables, so that a maximal distance travelled by odor molecules during the whole inhalation (*t*_*inh*_) averaged across all trials is equal to 1: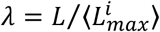. Here <…> means averaging across all trails,

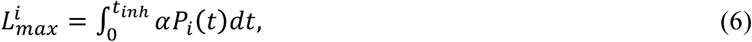

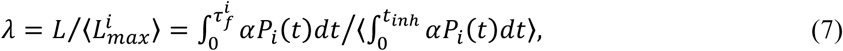

Thus, the relationship between *λ*, 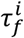, and *P*_*i*_(*t*) is as follows:

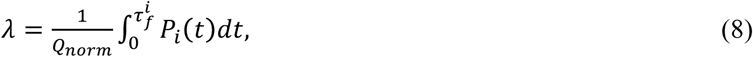

where

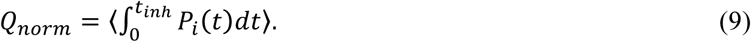

We first using equation (9), estimate *Q*_*norm*_, and normalize all pressure traces (Fig. 4B,C). Then, for a fixed λ, at a given trial, we estimate fluid dynamic delay time, 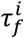, using equation (8), and shift spiking activity for a time interval 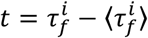, where 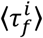> is an average fluid dynamic delay across all trials (Fig. 4C, D, and S2). Using shifted trails, we estimate PSTH and an average log-likelihood of individual spike trains fit by an average PSTH on the held out data. We varied λ between 0 and 1, with a step size of 0.05. Estimation of λ_*opt*_ that provides the best fit, was assessed by a polynomial fit of the second order to the dependence of log-likelihood on λ. We assumed that the best fit corresponded to minimal neuronal variability due to the sniff waveform signal.

<u>Model comparison.</u> To fit each model, we transform the spike trains 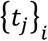 in individual trial *i* according to that model’s alignment rule (Fig. 5A, S4):

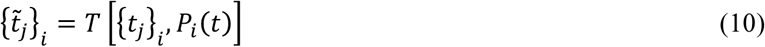

where *P*_*i*_(*t*) is the sniff waveform in trial *i* and *T* is an alignment rule. Then we calculated PSTHs of transformed spike trains:

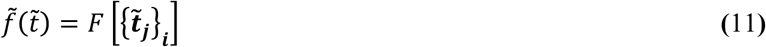

To evaluate the goodness of fit of different models, we evaluate their average log-likelihood in held out sniff cycles (test set). To this end, we first inversely transform each model’s PSTH according to the duration parameters of a given sniff cycle in the test set using interpolation.

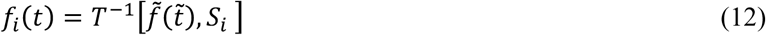

We then evaluate the Poisson process log-likelihood, given the inversely transformed PSTH, for the spike train during that sniff cycle (Fig. S5B,D):

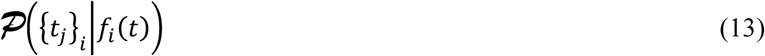

Finally, for each model we average the calculated single-sniff log-likelihoods over all sniffs to obtain that model’s average log-likelihood. We performed this analysis on 192 excitatory responses each consisting of at least 30 trials per response.

<u>Odor concentrations classification analysis.</u> For each concentration, we calculate PSTHs in real time and after FD alignment using the methods described in the Model comparison section above. For each alignment type, we discriminate between low and high concentrations using a likelihood ratio test. To this end, for each concentration, we evaluate the log-likelihood of a randomly drawn single trial spike train given the inversely transformed PSTH for that concentration. Repeating this procedure 300 times, we calculate the probability that the log-likelihood given the PSTH of the correct concentration is larger than log-likelihood given the PSTH of the other concentration. This yields a measure of discrimination success rate.

**Figure S1.**
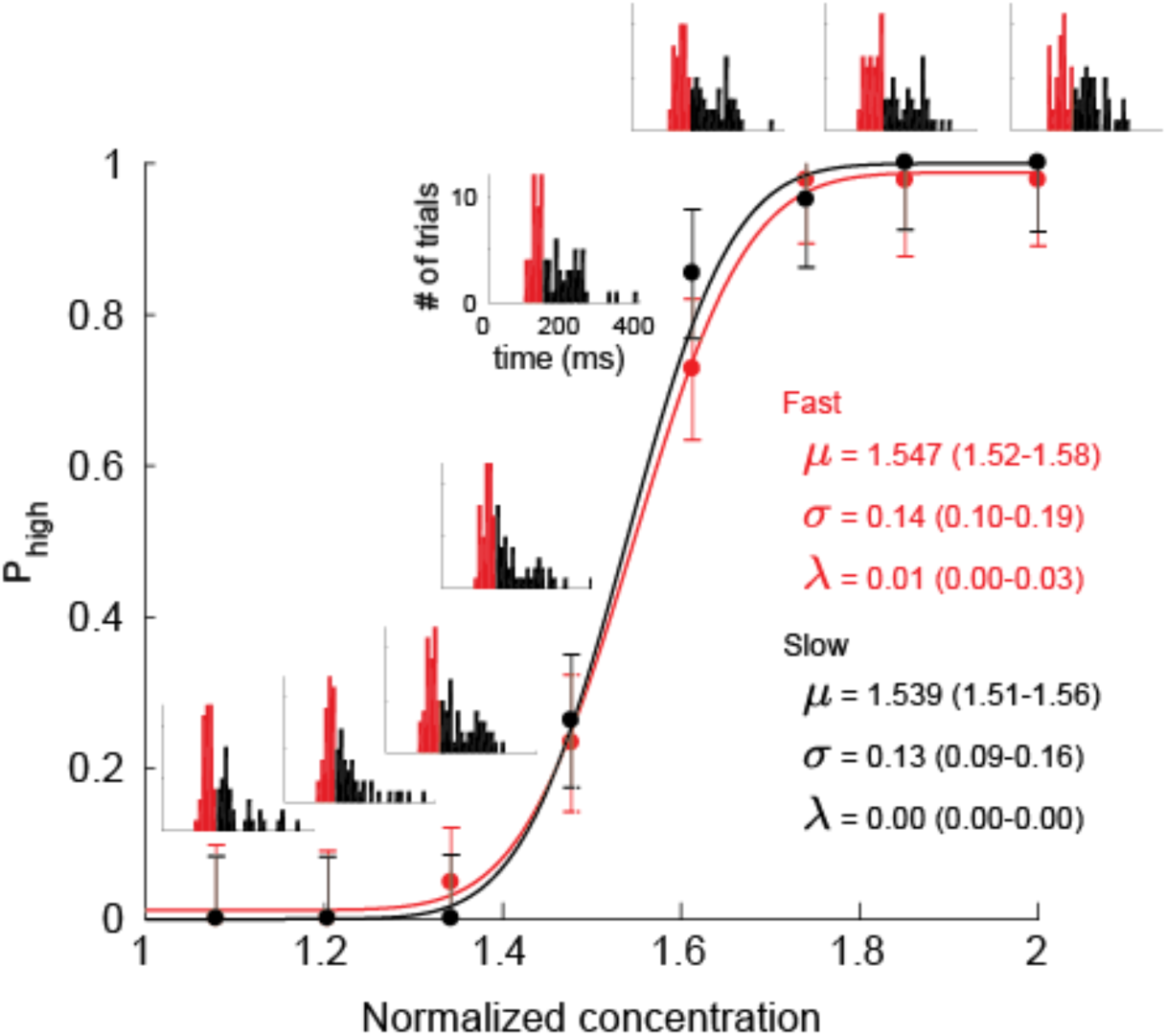
Example of a behavioral performance during a single session. Related to Figure 1. Performance during a single session for fast (black) and slow (red) sniff cycles. Each point denotes probability of going left (high concentration). Each inset is a sniff duration histogram for a given concentration.

**Figure S2.**
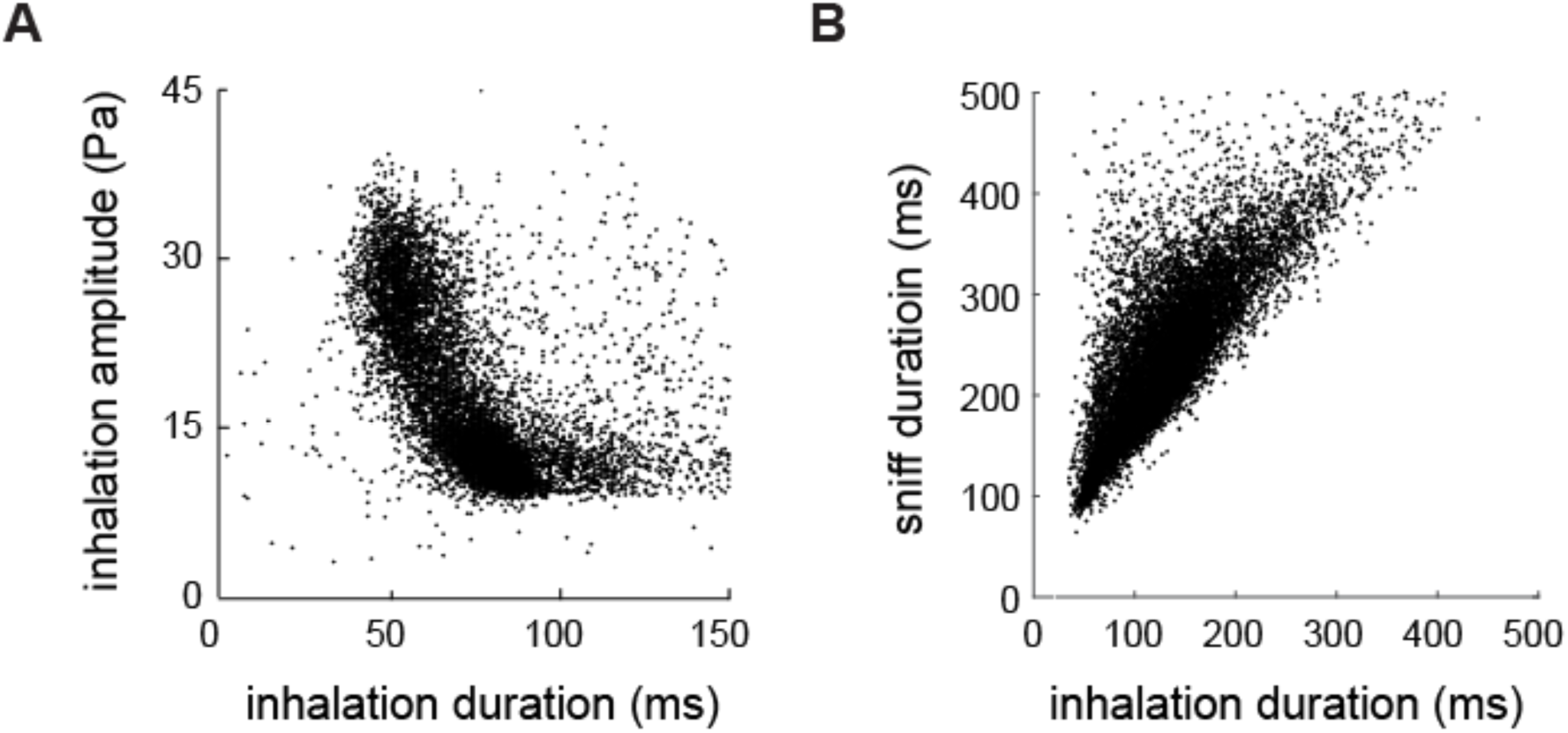
Comparisons of sniffing parameters. A. Comparison scatter plot of inhalation duration vs. inhalation amplitude. **B.** Scatter plot of inhalation duration vs. sniff durations, for a single mouse.

**Figure S3.**
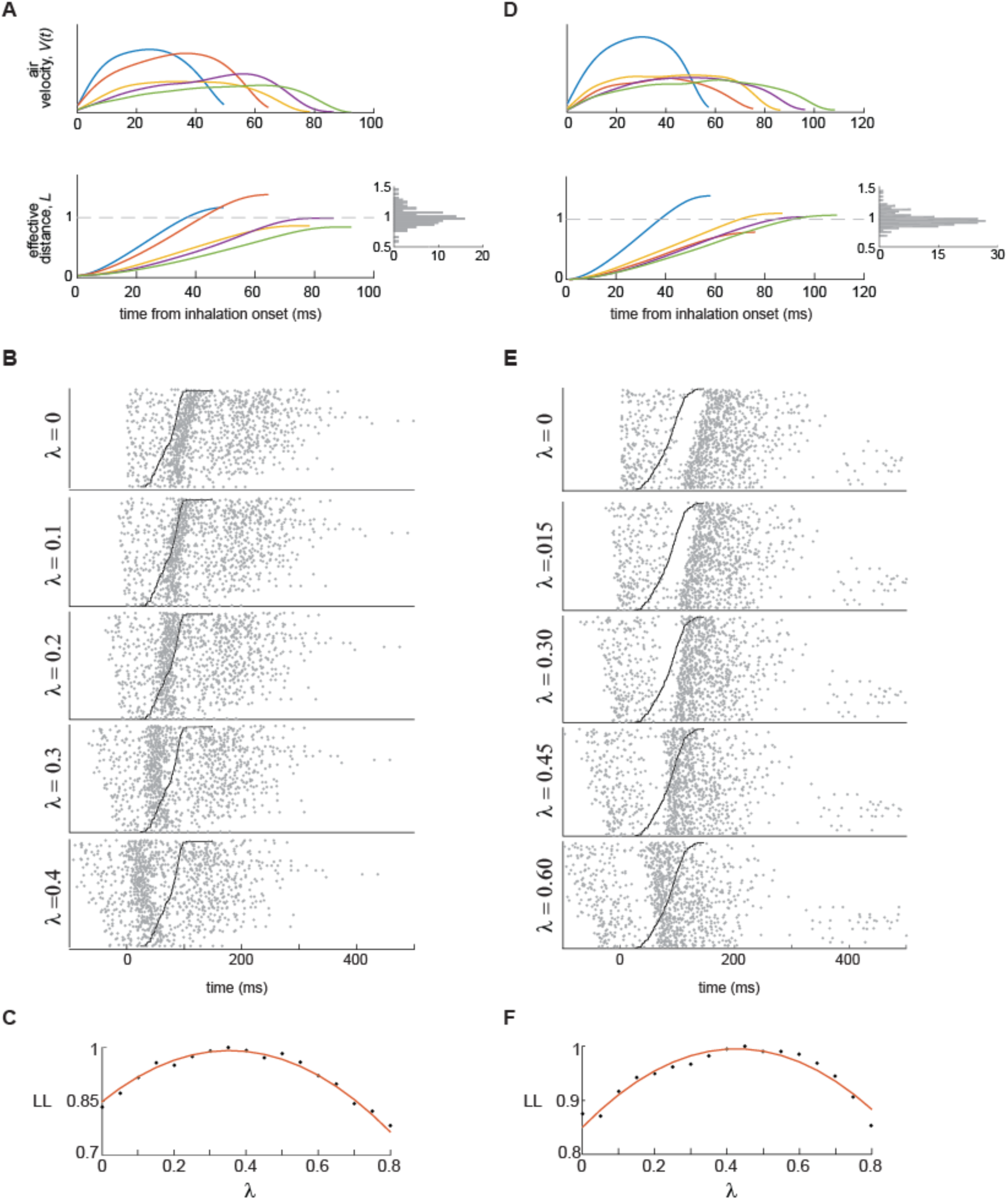
Examples of fluid dynamics model alignment. Related to Figure 4. **A.** Example of sniff waveforms and corresponding integrals. Right panels: a histogram of distribution of integrals over an inhalation interval. **B.** Raster plots of the first odorized sniff cycle, aligned to five different values of λ. Trials in the raster plots are ordered bottom to top by increasing inhalation duration. Black line shows end of the inhalation. **C.** Normalized log-likelihood (LL) as function of fraction of inhaled volume. Red curve is a parabolic fit to estimate optimal λ **D.-F.** same as **A.-C.** for different cell-odor pair.

**Figure S4.**
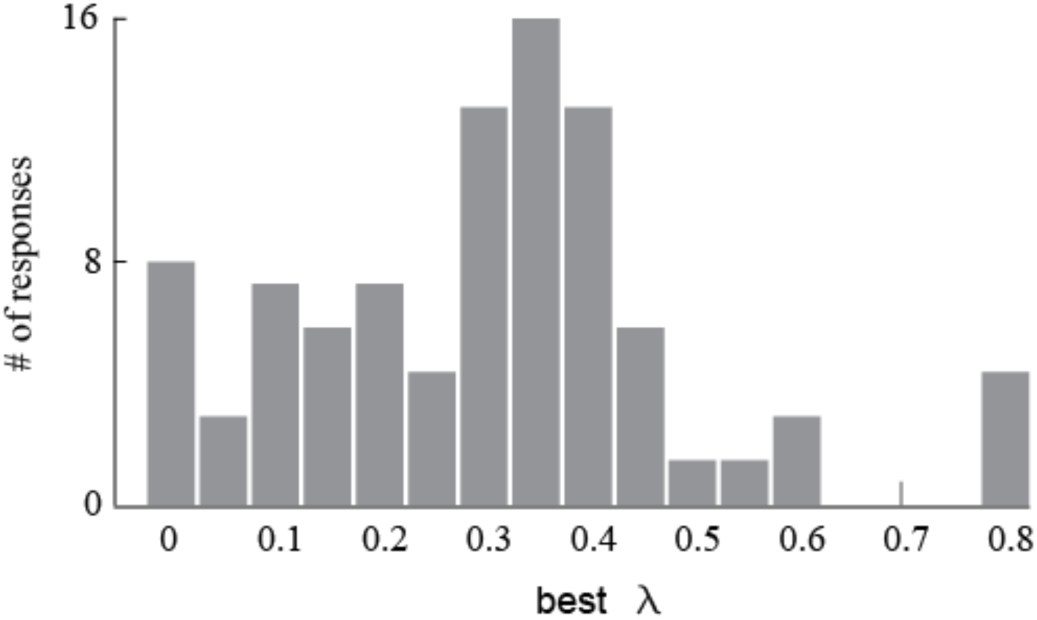
Distribution of the best λ parameters. Related to Figure 4. A histogram of distribution of λs for different cell-odor pairs.

**Figure S5.**
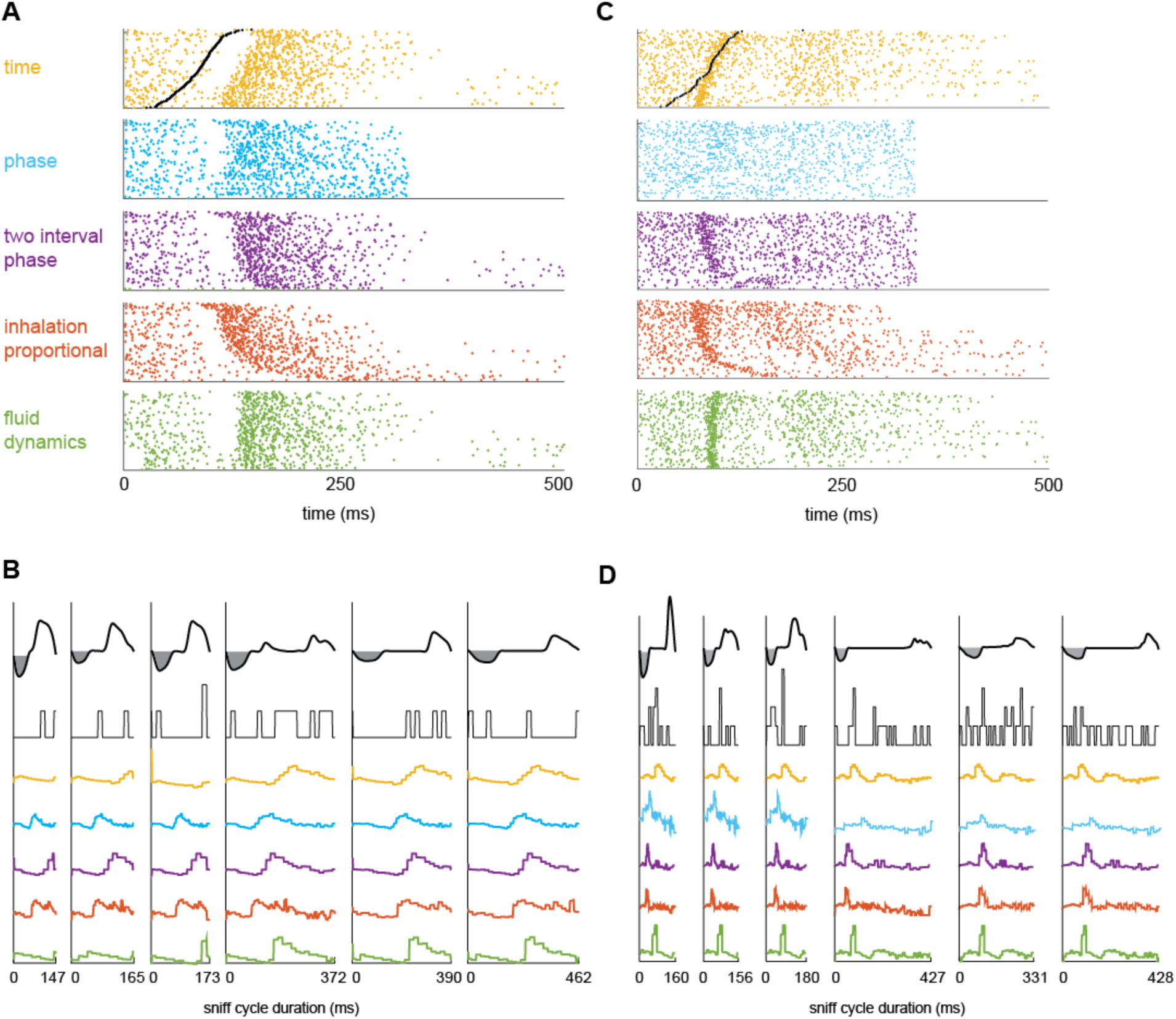
Performance of different alignment models. Related to Figure 5. **A.** and **C.** Raster plot of one cell-odor pair’s response, aligned according to different alignment models: Aligned by time (yellow); by phase, in which each sniff is warped to the average sniff duration (blue); two interval phase, in which each inhalation interval and the reminder of the sniff durations are equal to their average values (purple); inhalation proportional, in which inhalation is warped and the rest of the sniff is in real time coordinated (red); Fluid dynamic model, in which responses are aligned to the optimal effective length (λ) **B.** and **D.** Sniff waveforms, instantaneous firing rate from a single trial, and reconstructed PSTH plots from 5 models for 6 different sniff cycles.

**Figure S6.**
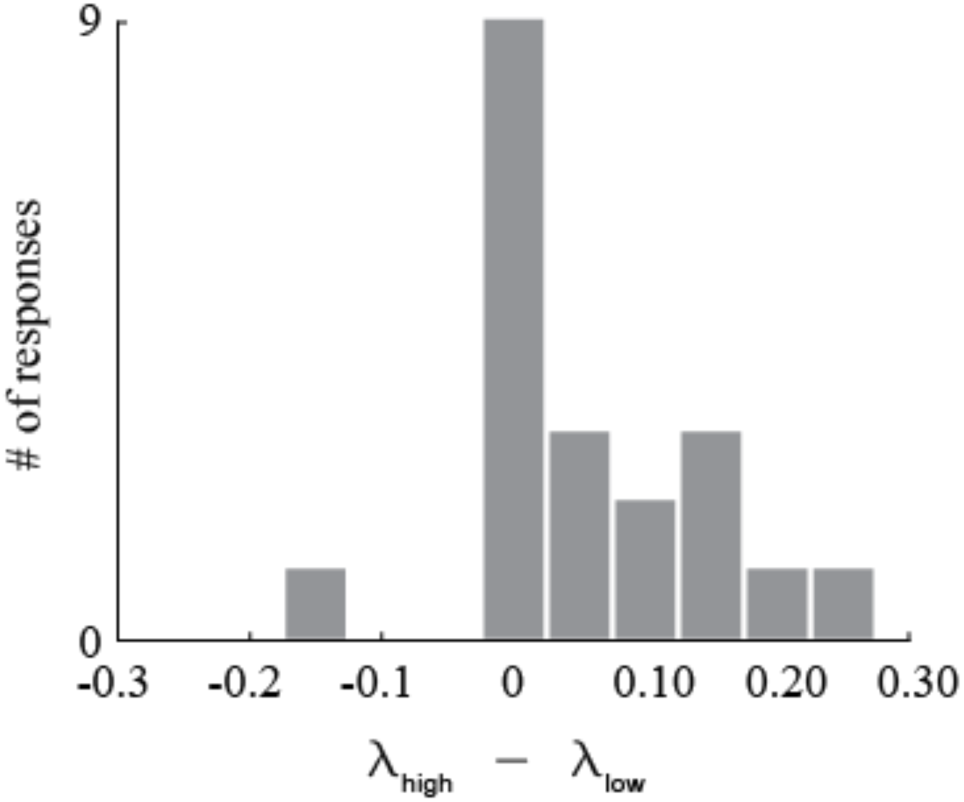
Related to Figure 7. Differences in λ between high and low concentrations (three-fold difference) for the same cell-odor pair. (Wilcoxon signed rank test, p=0.02)

